# *Legionella pneumophila* inhibits type I interferon signaling to avoid cell-intrinsic host cell defense

**DOI:** 10.1101/2022.12.21.521455

**Authors:** C.N.S. Allen, D. A. Banks, M. Shuster, S. N. Vogel, T.J. O’Connor, V. Briken

## Abstract

The host type I interferon (IFN) response protects against *Legionella pneumophila* infections. Other bacterial pathogens inhibit type I IFN-mediated cell signaling; however, the interaction between this signaling pathway and *L. pneumophila* has not been well described. Here, we demonstrate that *L. pneumophila* inhibits the IFN-β signaling pathway but does not inhibit IFN-γ-mediated cell signaling. The addition of IFN-β to *L. pneumophila*-infected macrophages limited bacterial growth independently of NOS2 and reactive nitrogen species. The type IV secretion system of *L. pneumophila* is required to inhibit IFN-β-mediated cell signaling. Finally, we show that the inhibition of the IFN-β signaling pathway occurs downstream of STAT1 and STAT2 phosphorylation. In conclusion, our findings describe a novel host cell signaling pathway inhibited by *L. pneumophila* via its type IV secretion system.

## Introduction

*Legionella* species are ubiquitous in the environment and have adapted to survive within freshwater amoebas (Rowbotham, 1980; Boamah et al., 2017). The bacteria rely on a type IV secretion system (T4SS) named defective in organelle trafficking and intracellular multiplication (Dot/Icm) to inject proteins into the host cell to manipulate host response pathways and increase their chances of survival (Berger and Isberg, 1993; Brand et al., 1994; Segal and Shuman, 1998; Vogel et al., 1998). *Legionella* can infect a wide variety of amoeba species and, therefore express multiple homologs of bacterial effectors to successfully modify the same host cell target in various hosts (Park et al., 2020). Humans are accidental hosts that can be exposed to *Legionella* via the aerosolization of contaminated water supplies, which may result in respiratory infections (McDade et al., 1977).

Type I interferons (IFNs), such as IFN-β, bind to the heterodimeric interferon-α/β receptor (IFNAR) and induce cell signaling, leading to the upregulation of hundreds of IFN-stimulated genes (ISGs) (Stetson and Medzhitov, 2006; Lazear et al., 2019). Type I IFNs are critical for host defense against viral pathogens (Stetson and Medzhitov, 2006; Lazear et al., 2019). However, during bacterial infections the activity of type I IFNs is more complex, in some cases mediating host resistance but in other cases increasing host susceptibility (Boxx and Cheng, 2016; Peignier and Parker, 2021). *Legionella pneumophila* induces IFN-β production by host cells via various cytosolic surveillance pathways, including the nucleic acid surveillance pathway (Liu and Shin, 2019). Type I IFNs then signal through IFNARs expressed on the cell surface of infected and bystander cells, contributing to *L. pneumophila* resistance, which has been observed in mice (Lippmann et al., 2011). Multiple studies have demonstrated that cell-intrinsic host defense mechanisms, including the production of nitric oxide (NO) (Plumlee et al., 2009) and itaconic acid (Naujoks et al., 2016), are triggered by exogenous IFN-β and subsequent IFNAR signaling during *ex vivo L. pneumophila* infections (Schiavoni et al., 2004; Opitz et al., 2006; Plumlee et al., 2009; Majoros et al., 2016). Thus, the IFN-β response plays a role in determining the outcome of *L. pneumophila* infections.

Here, we demonstrate that *L. pneumophila* can inhibit IFN-β-mediated cell signaling, and this inhibition occurs downstream of the phosphorylation of STAT1 and STAT2. Additionally, we show that the ability to reduce IFN-β signaling depends on the T4SS of *L. pneumophila* and is thus likely mediated by a secreted effector(s).

## Materials and Methods

### Cell culture and Mice

RAW 264.7 and RAW-Lucia ISG-KO-IRF3 cells (InvivoGen, rawl-koirf3) were grown in DMEM supplemented with 10% heat-inactivated fetal bovine serum (FBS). RAW-Lucia ISG-KO-IRF3 cells were plated in FBS-free DMEM 16 hours before the infection since the absence of FBS reduces the background activation of the integrated ISRE-Luciferase gene cassette.

THP-1 cells were purchased from ATCC and were maintained at 37°C and 5% CO_2_ in RPMI (ATCC) supplemented with 10% heat-inactivated FBS. For differentiation, THP-1 cells were treated with 50 ng/ml of phorbol 12-myristate 13-acetate (PMA) for two days. The media was then replaced and cells were cultured for two more days before infections.

Bone marrow-derived macrophages (BMDMs) were prepared from bone marrow cells flushed from the tibias and femurs of wild-type, *Ifnb*^-/-^, and *Nos2^-/-^* mice. The BMDMs were cultured in Gibco Dulbecco’s Modified Eagle Medium (DMEM) supplemented with 10% heat-inactivated FBS, 1% penicillin/streptomycin, and 20% L929 conditioned medium for six days prior to infection. L929 conditioned medium was used for all the BMDM infection experiments. *Ifnb*^-/-^ mice were originally obtained from Dr. E. Fish (University of Toronto) and have been previously described (Deonarain et al., 2000). 57BL/6J (JAX SO: 5530121) and *Nos2*^−/−^ (JAX SO: 5530126) mice were obtained from the Jackson Laboratory. All mice used for BMDMs were female.

### Ethics statement

All animals were handled in accordance with the NIH guidelines for the housing and care of laboratory animals. Studies were approved by the Institutional Animal Care and Use Committee at the University of Maryland.

### Bacteria

*L. pneumophila* Philadelphia-1 strain Lp02 was cultured at 37°C in liquid N-(2-87Acetamido)-2-aminoethanesulfonic acid (ACES)-buffered yeast extract (AYE) medium or on solid charcoal ACES-buffered yeast extract (CYET) containing 0.135 mg/ml L-cysteine (Sigma), 0.4 mg/ml iron(III) nitrate (Sigma), and 0.1 mg/ml thymidine (Sigma). The *L. pneumophila* Lp02 *dotA deficient* mutant (*ΔdotA)* was previously described (Berger and Isberg, 1993).

Heat inactivation of *L. pneumophila and ΔdotA L. pneumophila* was achieved based on a previous protocol (Allegra et al., 2011). Briefly, the bacteria were incubated in a water bath at 70°C for 30 minutes prior to the infections.

### Cellular infections

*L. pneumophila* were grown in AYE medium to the post-exponential phase (OD_600_ = 3.7– 4.1). RAW-Lucia ISG-KO-IRF3 cells, differentiated THP-1 cells, and BMDMs were infected with *L. pneumophila* for one hour at the indicated multiplicity of infections (MOI). The cells were washed three times with phosphate-buffered saline (PBS) and incubated in DMEM media supplemented with or without IFN-β for the given time.

### IFN-β enzyme-linked immunosorbent assay (ELISA)

The levels of secreted IFN-β were measured by an IFN-β enzyme-linked immunosorbent assay (ELISA) (R&D Systems) following the manufacturer’s protocol. RAW 264.7 cells were infected with *L. pneumophila* at an MOI of 0.05 for one hour. After one hour, the media was removed, the cells were rinsed three times with growth media, and fresh growth media was added. The supernatants of the infected and uninfected cells were collected at 24-, 48-, 72-, and 96-hours post-infection (h.p.i). IFN-β levels were measured by a plate reader (BioTek ELx800).

### Luciferase Reporter Assay

RAW-Lucia ISG-KO-IRF3 cells were infected with *L. pneumophila* at an MOI of 10 for one hour. After one hour, the cells were rinsed three times with media and then incubated in fresh growth media. At the indicated concentrations, IFN-β (R&D Systems) or IFN-γ (R&D Systems) was added to the post-infection media at 0 h.p.i. The supernatant from the infected and uninfected cells was collected, and the amount of type I IFN signaling was indirectly measured using QUANTI-Luc Gold, following the manufacturer’s protocol.

### qRT-PCR Assay

Total RNA was extracted using a GeneJet RNA Purification Kit (Thermo Scientific). A NanoDrop 8000 spectrophotometer (Thermo Scientific) was used to determine the purity and concentration of the RNA. cDNA was synthesized from collected RNA using SuperScript IV VILO cDNA Master Mix with ezDNase (Invitrogen). The following primers were used: BST2 (tetherin): (F) 5′-caaactcctgcaacctgaccgt −3′; (R) 5′-ctcctggttcagcttcgtgact-3′ and beta actin: (F) 5′-cattgctgacaggatgcagaagg-3′; (R) 5′-tgctggaaggtggacagtgagg-3′. All analyses were performed using FastStart Universal SYBR Green (Roche) according to the manufacturer’s instructions. The relative mRNA quantitation was done using the comparative ΔΔCt method, and all results are compared to the control group and loading control beta actin.

### Cell Death AK Release Assay

A ToxiLight BioAssay Kit (Lonza) was used to determine cell death. This assay measures the amount of adenylate kinase (AK) in the cellular supernatant released upon cell membrane damage. The assay was performed following the manufacturer’s protocol. Briefly, 20 μl of cellular supernatants from the infected and uninfected cell cultures were added to a 96-well plate. Then, 100 μl of the AKDR solution from the kit was added to each well and incubated for five minutes. The plate was then read using a microplate reader (BioTek).

### Bacterial Survival

For bacterial survival assays, *L. pneumophila* used for infections was grown to the post-exponential phase (OD_600_ = 3.8–4.0). Wild-type, *Nos2^-/-^*, and *Ifna^-/-^* BMDMs were infected with *L. pneumophila* at an MOI of 1 for one hour. After one hour, the cells were rinsed three times, and the media was replaced with BMDM growth media. At the provided time points, saponin was added to the cells to achieve a final concentration of 0.02% (w/v), and the cells were incubated for ten minutes at 37°C. The medium was then harvested, and 100 μL of sterile water was added to the wells. The water was then harvested and mixed with the original harvested media. The bacteria were counted based on colony-forming units (CFUs) from plating diluted cellular lysates on CYE medium agar plates.

### Antibody neutralization

Antibody neutralization studies were performed using *InVivo*Plus anti-mouse IFNAR1 and *InVivo*MAB mouse IgG1 isotype control antibodies (Bio X Cell). Antibodies were added to the infected cells at 0 h.p.i. at a final concentration of 20 μg/ml. The bacterial survival study was performed using wild-type 57BL/6J BMDMs. Briefly, BMDMs were infected with *L. pneumophila* at an MOI of 1 for one hour, then the cells were rinsed three times, and the media was replaced. At the provided time points, saponin was added to the cells to achieve a final concentration of 0.02% (w/v), and the cells were incubated for ten minutes at 37°C. The medium was then harvested, and 100 μL of sterile water was added, harvested, and mixed with the original harvested media. The bacteria were counted based on colony-forming units (CFUs) from plating diluted cellular lysates on CYE medium agar plates.

### NO assay

The production of NO in response to IFN-β was quantified using a Griess assay kit (Promega) following the manufacturer’s protocol. Briefly, 50 μL of supernatant from the infected cells with or without IFN-β at the indicated time points was added to wells in triplicate, mixed with 50 μL of the Sulfanilamide solution, and incubated at room temperature for ten minutes. Then, 50 μL of the NED solution was added and incubated at room temperature for ten minutes. The production of nitrite, a derivative of NO, was measured at 548 nm using a microplate reader (BioTek). The nitrite concentration was determined using a sodium nitrite standard curve.

### Flow Cytometry

Differentiated THP-1 cells were infected with GFP-positive *L. pneumophila or L. pneumophila ΔdotA* at an MOI of 10 for one hour. After infection, the media was removed, cells were rinsed three times with fresh media, and fresh growth media was added and incubated for 30 minutes (STAT1 and STAT2) or 20 hours (tetherin). At collection, the medium was removed, and Cellstripper (Corning) was added to detach the cells from the plate surface. The cells were centrifuged at 425 x g for three minutes at 4°C. The supernatant was removed, and the cell pellet was rinsed twice with 1 ml of PBS.

For live/dead staining, a LIVE/DEAD Yellow Fixable Dead Stain kit (Thermo Fisher Scientific) was used for the anti-tetherin flow cytometry, and a LIVE/DEAD Near-IR Fixable Dead Stain kit (Thermo Fisher Scientific) was used for the STAT1 and STAT2 flow cytometry following the manufacturer’s protocol. Briefly, the collected and rinsed cell pellets were resuspended in 1 ml of PBS, and 1 μL of the fluorescent reactive dye from the kit was added to each sample. The cells were then incubated in the dark for 30 minutes. The cells were then centrifuged at 425 x g for three minutes at 4°C, rinsed with 1 ml of PBS, and centrifuged two additional times. After the last centrifugation, the cell pellets were resuspended in the flow buffer (PBS with 0.5 % BSA).

For the cell surface staining for the tetherin flow cytometry, the cell pellets were resuspended in 50 μL of flow buffer. Human TruStain FcX (BioLegend) was added at a 1:10 dilution, and the cells were briefly vortexed. The cells were incubated for ten minutes. Next, 50 μL of the stain cocktail (1x Brillant Stain Buffer [BD Bioscience] and 5 μL per sample of APC anti-human CD317 (BST2, Tetherin) antibody [BioLegend]) was added to each sample. The samples were then incubated for one hour on ice in the dark. After the incubation, 1 ml of flow buffer was added to the samples, centrifuged at 425 x g for three minutes at 4°C, rinsed with 1 ml of flow buffer, and centrifuged two more times. The cells were resuspended in 4% PFA before running the samples.

For the intercellular staining for STAT1 and STAT2, the cell pellets were resuspended in 100 μL of 4% PFA for 15 minutes on ice. The cells were centrifuged at 425 x g for three minutes at 4°C and rinsed with ice-cold PBS. For permeabilization, 1 ml of ice-cold 100% methanol was added to the cells on ice. The cells were incubated for ten minutes on ice. The cells were then centrifuged at 425 x g for three minutes at 4°C and rinsed with ice-cold PBS. The cell pellets were resuspended in 50 μL of flow buffer. Human TruStain FcX (BioLegend) was added at a 1:10 dilution, and the cells were briefly vortexed. The cells were incubated for ten minutes. Next, 50 μL of the stain cocktail (1x Brillant Stain Buffer [BD Bioscience] and either 5 μL per sample of Alexa Flour 647 Anti-Total Stat1 [N-Terminus] [BD Biosciences] and 5 μL per sample of PE anti-STAT1 Phospho [Tyr701] [BioLegend]; or 2 μL per sample of Stat2 [D9J7L] Rabbit mAb [PE] [Cell Signaling Technology] and 2 μL per sample of P-Stat2 [P-690] Rabbit mAb [Alexa 647] [Cell Signaling Technology]) was added to each sample. The samples were then incubated for one hour on ice in the dark. After the incubation, 1 ml of flow buffer was added to the samples, centrifuged at 425 x g for three minutes at 4°C, rinsed with 1 ml of flow buffer, and centrifuged two more times. The cells were then resuspended in 4% PFA prior to running the samples.

### Statistical Analysis

All statistical analyses were performed using GraphPad 9.0. For all the experiments, the data are shown as the mean ± standard deviation (*SD*) of three biological replicates unless otherwise noted. Significance is indicated by a *p-*value less than or equal to 0.05.

## Results

To test how much IFN-β is produced during *L. pneumophila* infections, RAW 264.7 cells were infected with *L. pneumophila* at an MOI of 0.05 for one hour. After one hour of infection, the cells were rinsed, and fresh media was added. The amount of secreted IFN-β was measured at 24-hour intervals via ELISA. Each data point is obtained from the average of three technical replicates and represents an independent experiment. After 24 hours, 163.5 ± 55.2 pg/ml of IFN-β was detected, a ten-fold increase compared with the uninfected cells, and the IFN-β production stayed constant until 48 h.p.i. At 72 h.p.i., the level of IFN-β increased to 342.4 ± 103.9 pg/ml and continued to rise to 460.9 ± 109.5 pg/ml by 96 h.p.i. (Figure 1A). These results indicate that *L. pneumophila* induces IFN-β production in RAW 264.7 macrophages and that IFN-β accumulates over time after the infection.

**Figure 1:**
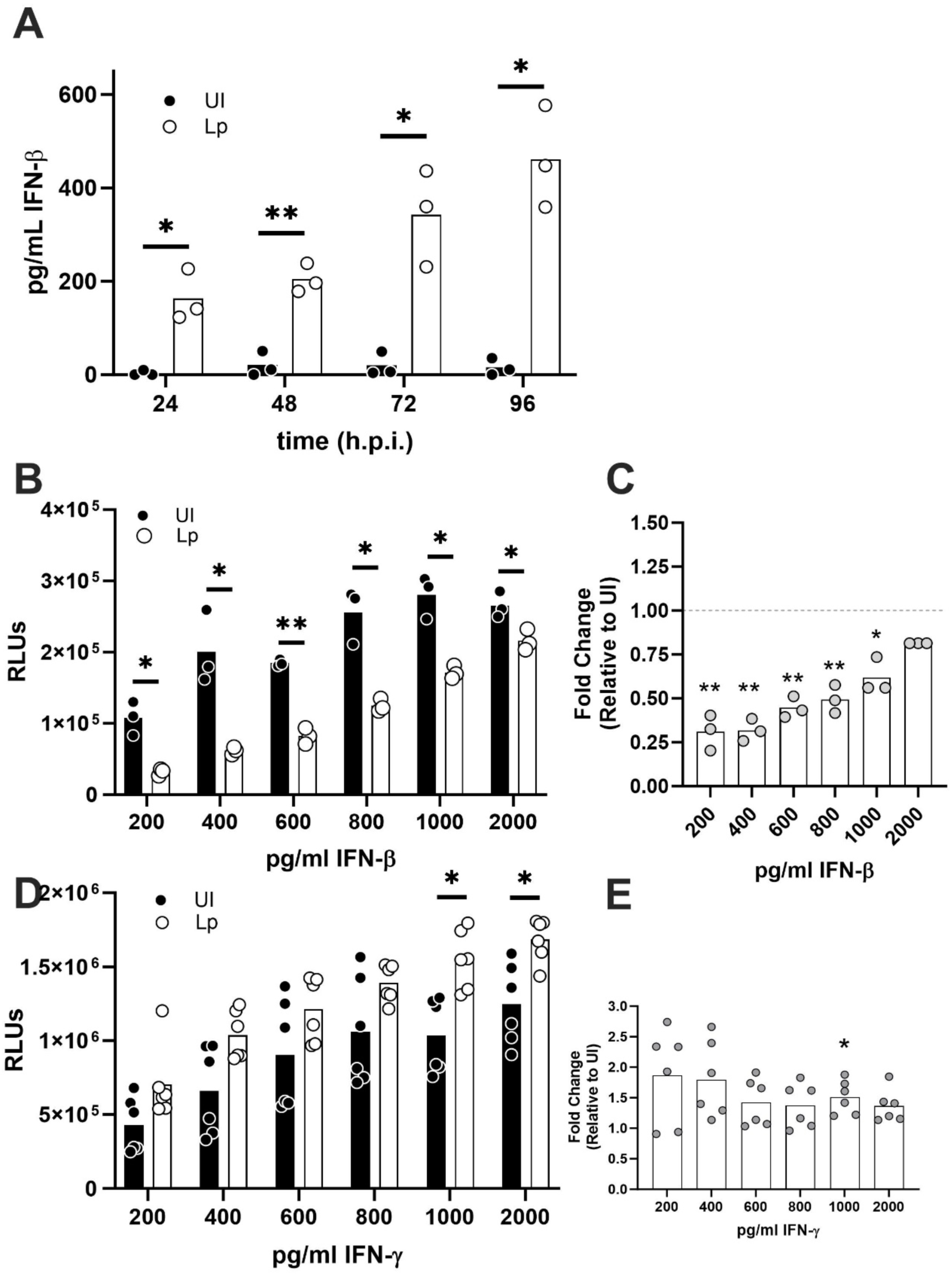
*Legionella pneumophila* inhibits type I but not type II IFN signaling. (A) RAW-Lucia ISG-KO-IRF3 cells were infected with *L. pneumophila* (Lp) at an MOI of 0.05, and the amount of secreted IFN-β was detected by ELISA. (B) Infected (Lp) (MOI of 10) and uninfected (UI) cells were challenged with increasing concentrations of IFN-β, and the IFN-β/γ-signaling reporter gene expression was measured after 20 hours post-infection (h.p.i.) by detection of secreted luciferase into the supernatant and values are plotted as relative light units (RLU). (C) The data in B was expressed as the fold change of Lp-infected, IFN-β-treated cells compared to uninfected, IFN-β-treated cells. (D) Infected (MOI of 10) and uninfected cells were challenged with increasing concentrations of IFN-γ, and the IFN-β/γ-signaling reporter gene expression was measured after 20 h.p.i. as for B. (E) The data in D was expressed as the fold change of Lp-infected, IFN-γ-treated cells compared to uninfected, IFN-γ-treated cells. Individual points are from independent experiments and indicate the averages of three technical replicates. Statistical analyses were done using multiple unpaired *t-*tests with Welch correction for each time point (A) or each dose concentration of IFN-β or IFN-γ (B, D). The fold change is relative to the uninfected cells for the same IFN-β or IFN-γ treatment concentration. Statistical analyses were done using multiple unpaired *t*-tests with Welch correction, and each fold change was compared to the fold change of the uninfected cells for each treatment dose of IFN-β or IFN-γ (C, E). Significance is indicated as follows: \**p* ≤ 0.05 and \*\**p* ≤ 0.01.

Since it has been previously reported that *Mycobacterium tuberculosis* and *Shigella sonnei* inhibit IFNAR signaling in infected cells (Banks et al., 2019a; Alphonse et al., 2022), we wanted to determine if *L. pneumophila* also inhibits type I interferon signaling. RAW-Lucia ISG-KO-IRF3 cells were used for these studies since they are deficient in *Irf3* and consequently infections with *L. pneumophilia* will not induce the production of endogenous IFN-β. Hence, the only IFN-β-mediated cell signaling detected by the IFN-β-inducible reporter gene is that of the IFN-β added exogenously to the cells. Cells infected or not with *L. pneumophila* at an MOI of 10 were cultured with increasing amounts of IFN-β (200–2000 pg/ml) for 20 hours to measure the effect *L. pneumophila* has on IFN-β-mediated cell signaling. The secretion of luciferase into the supernatant correlates with the activity of the IFN-β responsive promotor. The secreted luciferase was measured via a chemiluminescence assay and results are expressed in relative light units (RLU). Cells infected with *L. pneumophila* had lower levels of IFN-β-associated gene expression at all the doses of IFN-β tested when compared to the uninfected, untreated cells (Figure 1B). At the lower doses of IFN-β (200 and 400 pg/ml) a fold change of 0.25 relative to the uninfected cells was measured which changed to 0.8 at the highest IFN-β dose of 2000 pg/ml (Figure 1C). The IFN-β signaling pathway inhibition was reduced in a dose-dependent manner, suggesting that as the IFN-β concentration increases, the ability of *L. pneumophila* to impede the signaling pathway is overwhelmed.

The IFN-β and IFN-γ signal transduction pathways share some of their intracellular signaling components, and thus, we asked if *L. pneumophila* can also inhibit IFN-γ signaling. The RAW-Lucia ISG-KO-IRF3 reporter construct is also induced by IFN-γ signaling and could thus serve as a model to study the potential of *L. pneumophila* to inhibit IFN-γ signaling. Increasing amounts of IFN-γ (200–2000 pg/ml) were added to *L. pneumophila*-infected or uninfected cells (MOI of 10), and the luciferase production was analyzed (Figures 1D and E). Overall, there was no reduction in the IFN-γ response in the infected cells; however, at IFN-γ doses of 1000 and 2000 pg/ml, there was a significant increase in transcription response in the *L. pneumophila*-infected cells compared with the uninfected cells (Figures 1D and E). These results indicate that *L. pneumophila* partially inhibits IFN-β signaling while not affecting IFN-γ signaling. *L. pneumophilia* can reduce host protein synthesis via the secretion of multiple effectors (Belyi et al., 2006; Shen et al., 2009; Fontana et al., 2011; Moss et al., 2019). To determine if the decrease in the IFN-β signaling in *L. pneumophila-*infected RAW-Lucia ISG-KO-IRF3 cells was due to decreased global protein production, a q-RT-PCR for the mRNA of the IFN-β-inducible gene tetherin (BST-2) was performed. In RAW-Lucia ISG-KO-IRF3 cells infected or not with *L. pneumophila* at an MOI of 10, treated or not with IFN-β and cells were harvested for mRNA extraction at 4 hours post infection. The level of tetherin mRNA in the infected cells, not treated with IFN-β was three-fold higher than in the uninfected RAW-Lucia ISG-KO-IRF3 not treated with IFN-β. The induction of tetherin mRNA in the infected RAW-Lucia ISG-KO-IRF3 cells treated with IFN-β (1000 pg/ml) was significantly less than the induction of tetherin mRNA in the uninfected RAW-Lucia ISG-KO-IRF3 treated with IFN-β (Figure 2A). This result suggests that the decrease in IFN-β signaling pathway is due to an inhibition of an event upstream of transcription initiation. It also shows that the observed inhibition in the generation of luciferase in the reporter cells is not due to an inhibition of overall protein synthesis due to *L. pneumophila* infections.

**Figure 2:**
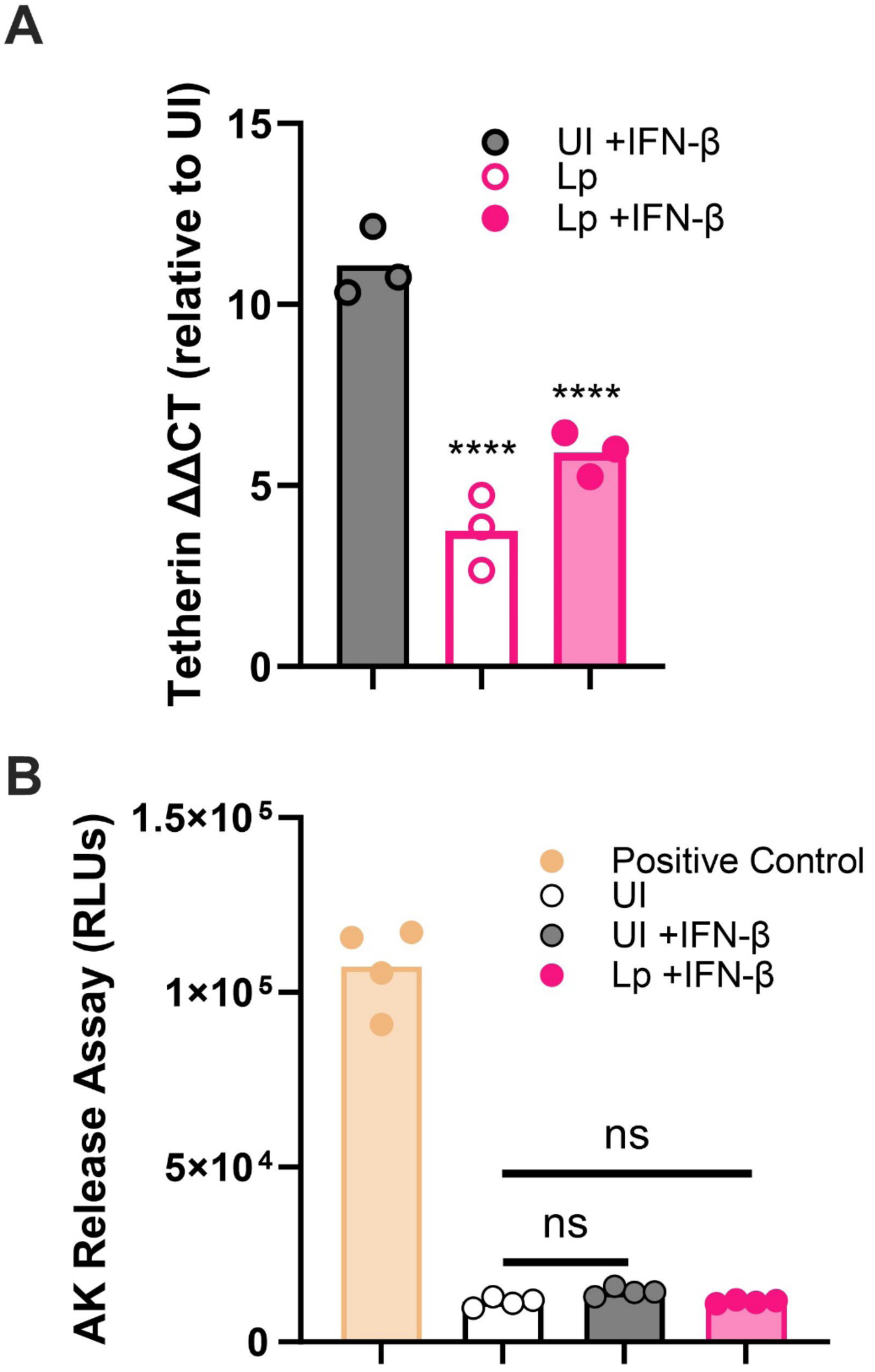
*L. pneumophila* (Lp) infection decreases transcriptional response to IFN-β-treatment without affecting cell death. (A) Tetherin mRNA expression of RAW-Lucia ISG-KO-IRF3 cells was quantified via qRT-PCR in *L. pneumophila* (Lp)-infected/untreated cells, Lp-infected/IFN-β-treated cells and uninfected (UI)/IFN-β-treated cells at 4 hours post infection (h.p.i.) and expressed as fold change compared to uninfected, untreated cells. The cells were infected with Lp at an MOI of 10. (B) RAW-Lucia ISG-KO-IRF3 cell death was measured by an adenylate kinase release assay in cells infected or not with *L. pneumophila* at an MOI of 10 after 20 h.p.i. and expressed as relative light units (RLU). The positive control is cells lysed with a detergent. For A and B, individual points are the averages of three technical repeats and obtained from independent experiments. An asterisk indicates multiple unpaired *t-*tests with Welch correction relative to the untreated cells. Significance is indicated as follows: \*\*\*\**p* ≤ 0.0001.

An AK release assay was performed to test if the decrease in the IFN-β-mediated luciferase production was due to a decrease in the total number of reporter cells due to increased cell death. After 20 h.p.i, compared with the uninfected, non-IFN-β treated RAW-Lucia ISG-KO-IRF3 cells, there was no significant increase in cell death in the *L. pneumophila*-infected (MOI of 10) IFN-β treated cells (Figure 2B). This result suggests that the decrease in luciferase in the *L. pneumophila*-infected RAW-Lucia ISG-KO-IRF3 cells was not from decreased cell counts via increased cell death.

IFN-β has been shown to contribute to the clearance of *L. pneumophila* in mice (Lippmann et al., 2011) and in *ex vivo* infected BMDMs (Schiavoni et al., 2004; Plumlee et al., 2009; Lippmann et al., 2011). *Ifnb^-/-^*-derived BMDMs were infected with *L. pneumophila* and treated with IFN-β to determine the effects of IFN-β on bacterial survival within macrophages. We used *Ifnb^-/-^*-derived BMDMs to observe the effects of exogenous IFN-β without the production of endogenous IFN-β in response to the *L. pneumophila* infection (Plumlee et al., 2009; Lippmann et al., 2011). Infected cells were treated with or without an IFNAR1-neutralizing antibody or an isotype antibody to demonstrate the specificity of the IFN-α treatment due to IFNAR signaling. The *Ifnb^-/-^*-derived BMDMs were infected with *L. pneumophila* at an MOI of 1, and then 20 μg/ml of the IFNAR1-neutralizing or isotype control antibodies were added. The IFN-β treated groups were then treated with 1000 pg/ml IFN-β. The addition of IFN-β to the *Ifnb^-/-^*-derived BMDMs did not affect the growth of the bacteria for the first two time points (24 and 48 h.p.i.) but at 72 and 96 h.p.i. there was a significant reduction in the CFUs in the isotype IFN-β-treated cells compared with the untreated isotype cells (Figure 3A). However, blocking the IFNAR receptor by the IFNAR1-neutralizing antibodies inhibited the IFN-β-mediated reduction in the CFUs (Figure 3A). These results indicate that IFN-β plays a role in the growth inhibition of *L. pneumophila* in macrophages and that this inhibition is dependent on signaling through the IFNAR1 receptor.

**Figure 3:**
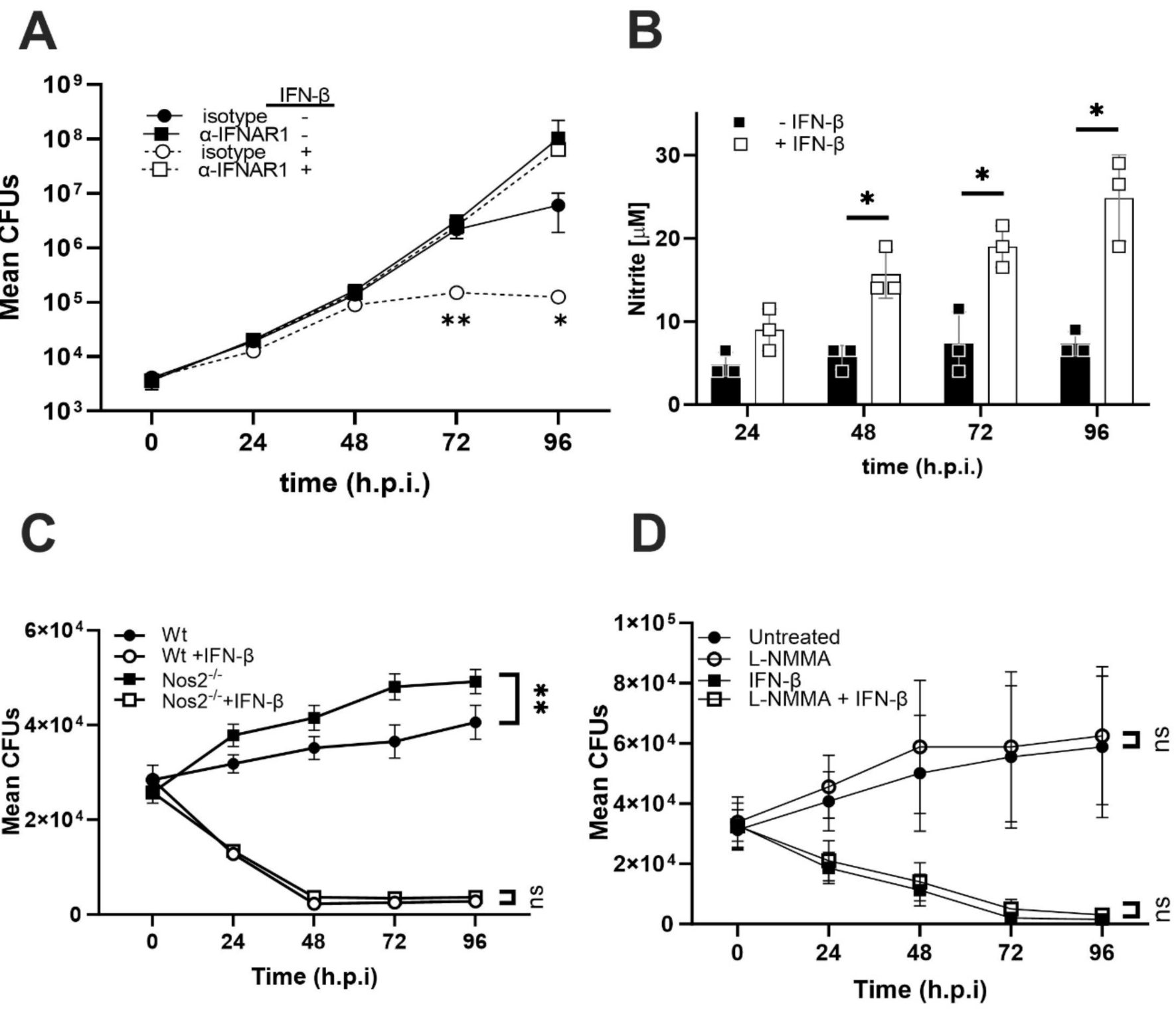
IFN-β-treatment increases cell-intrinsic resistance to *L. pneumophila* independent of NO production. (A) Bone marrow-derived macrophages (BMDMs) from *Ifnb^-/-^*mice were infected with *L. pneumophila* (Lp) at an MOI of 1 and treated with 1000 pg/ml IFN-β throughout the infection. The number of colony-forming units (CFUs) was counted at the indicated time points. The data are the mean ± standard deviation of six replicates obtained from two independent infections. Asterisks indicate multiple unpaired *t-*tests with Welch correction relative to the untreated cells. (B) RAW-ISG-KO-IRF3 cells were infected with Lp at an MOI of 10. The amount of produced nitrite was measured using a Griess assay and shown as the averages of three technical replicate from three independent experiments. (C) BMDMs from *Nos2^-/-^* mice were infected with Lp at an MOI of 1 and treated with 1000 pg/ml IFN-β throughout the infection. The number of CFUs was counted at the indicated time points. The points are the mean ± standard deviation of three independent experiments. Asterisks indicate multiple unpaired *t-*tests with Welch correction relative to the untreated cells. (D) Wildtype BMDM cells were infected with Lp at MOI of 1 and treated with 1000 pg/ml IFN-β and solvent control (DMSO) or L-NMMA (12 μM). The mean ± standard deviation of three independent experiments are shown. Asterisks indicate a multiple unpaired *t-*test with Welch correction comparing DMSO ± IFN-β or L-NMMA ± IFN-β. Significance is indicated as follows: \**p* ≤ 0.05 and \*\**p* ≤ 0.01.

Next, we investigated the role of NO production on host protection during IFN-α treatment. The NO production was analyzed using a Griess assay, which detects nitrite as a proxy for the induced production of NO. *L. pneumophila*-infected (MOI of 10) RAW-Lucia ISG-KO-IRF3 cells had a slight increase in nitrite levels in the IFN-β-treated cells (1000 pg/ml) compared with untreated cells at 24 h.p.i. (Figure 3B). After 48 hours, the IFN-β treatment increased the nitrite level to about 18 μM; however, the nitrite level in the untreated cells was about 5 μM. The IFN-β treatment led to nitrite production of about 25 μM at 96 hours, whereas the level in the untreated cells remained low and was around 8 μM (Figure 3B). These results show that IFN-β signaling induced NO production in the RAW-Lucia ISG-KO-IRF3 cells and that the production of NO is reduced in cells infected with *L. pneumophila*.

To determine if the reduced NO production influenced the clearance of *L. pneumophila,* wild-type and *Nos2^-/-^* BMDMs were infected with *L. pneumophila* at an MOI of 1 and treated with or without IFN-β (1000 pg/ml). The *Nos2^-/-^* BMDMs not treated with IFN-β had the greatest amount of bacterial growth. The wild-type IFN-β untreated BMDMs had significantly reduced bacterial growth over all the measured time points compared with the *Nos2^-/-^* untreated BMDMs (Figure 3C). This result suggests that Nos2 provides some protection against *L. pneumophila* in an IFN-β independent pathway. The *Nos2^-/-^* and wild-type BMDMs treated with IFN-β had significantly decreased bacterial survival compared with the BMDMs not treated with IFN-β; however, there was no significant difference between the *Nos2^-/-^* BMDMs and the wild-type BMDMs treated with IFN-β (Figure 3C). This lack of significance in bacterial survival between the *Nos2^-/-^*BMDMs and the wild-type BMDMs treated with IFN-β suggests that the growth inhibition of *L. pneumophila* observed with IFN-β is Nos2 independent.

To further test the role of reactive nitrogen species (RNS) in the clearance of *L. pneumophila* in response to IFN-β, the iNOS inhibitor L-NMMA was used. First, it was determined that L-NMMA had a dose-dependent effect on the nitrite production of wild-type BMDMs treated with IFN-β (1000 pg/ml) after 20 hours. At the highest dose of L-NMMA (12 μM), the nitrite production was approximately 15% of the nitrite produced by the untreated cells (Supplementary Figure 1A). L-NMMA (12 μM) was also determined to have no effect on *L. pneumophila* cell growth rate (Supplementary Figure 1B). Wild-type BMDMs were infected with *L. pneumophila* at an MOI of 1 and treated with or without IFN-β (1000 pg/ml) and with a solvent control (DMSO) or 12 μM of L-NMMA to observe the effect that NO production has on the survival of *L. pneumophila* (Figure 3D). The L-NMMA treated and IFN-β untreated BMDMs and the untreated BMDMS had similar bacterial growth at all the tested time points. However, the L-NMMA and IFN-β treated, and the only IFN-β treated BMDMs, had significantly decreased bacterial survival compared with the BMDMs not treated with IFN-β (Figure 3D). This decreased bacterial survival was similar for the L-NMMA and IFN-β treated and the only IFN-β treated BMDMs. This result further suggests that the clearance of *L. pneumophila* observed with IFN-β is independent of RNS. *L. pneumophila* T4SS Dot/Icm is a critical virulence factor that deploys effector proteins to the host cell to manipulate host cellular processes (Berger and Isberg, 1993; Brand et al., 1994; Segal and Shuman, 1998; Vogel et al., 1998). One mechanism by which *L. pneumophila* might inhibit IFN-β signaling is through the activity of a secreted effector. To test this, an *L. pneumophila dotA* deficient mutant (Δ*dotA*) was used to explore the importance of the Dot/Icm system and, thus, the possibility that an effector is involved in inhibiting the IFN-β signaling pathway. RAW-Lucia ISG-KO-IRF3 cells were infected with wild-type *L. pneumophila* or the Δ*dotA* mutant *L. pneumophila* at an MOI of 10 and were treated with 0, 600, 1000, or 2000 pg/ml of IFN-β for 20 hours (Figure 4A). The IFN-β-mediated signaling was reduced in the cells infected with the wild-type bacteria to approximately 60% of uninfected, IFN-β-treated cells, for all the tested IFN-β concentrations (Figure 4A). In contrast, the Δ*dotA* mutant did not reduce the IFN-β-mediated signaling as much as the wild-type *L. pneumophila* (Figure 4A). In addition to the wild-type and Δ*dotA* mutant, RAW-Lucia ISG-KO-IRF3 cells were infected with heat-inactivated wild-type or Δ*dotA* mutant bacteria at an MOI of 10 to explore if the inhibition of the IFN-β signaling pathway requires live bacteria or if the inhibition is a result of pre-existing heat resistant bacterial lipids. The two heat-inactivated bacterial strains did not inhibit IFN-β signaling compared with the uninfected Raw-Lucia ISG-KO-IRF3 cells (Figure 4A). Overall, these results suggest that the inhibition of the IFN-β pathway results from the activity of an effector secreted through the Dot/Icm T4SS.

**Figure 4:**
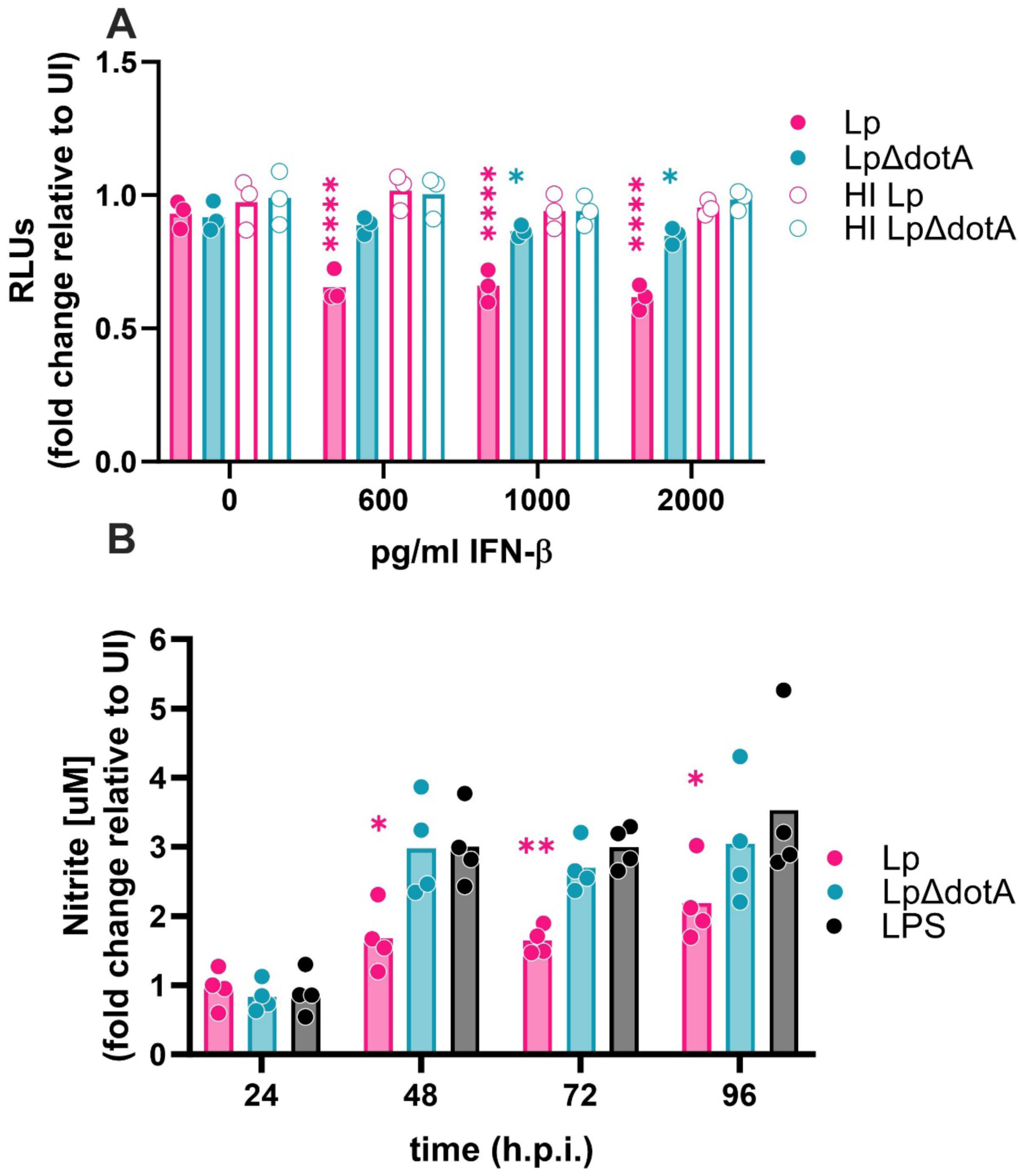
The inhibition of IFN-β signaling by *L. pneumophila* is dependent on its type IV secretion system. (A) RAW-ISG-KO-IRF3 cells were infected with wild-type *L. pneumophila* (Lp), the Lp *dotA* deficient mutant (LpΔ*dotA*), the heat-inactivated wild-type Lp (HI Lp), or the heat-inactivated *dotA* deficient mutant (HI LpΔ*dotA*) at an MOI of 10. All experimental groups were treated with the indicated amounts of IFN-β. The y-axis shows the fold change in IFN-β reporter gene expression compared to the uninfected/untreated experimental group after 20 h.p.i.. Data points are from independent experiments and represent the averages of three technical replicates. (B) RAW-ISG-KO-IRF3 cells were infected with wild-type Lp or the Lp *dotA* deficient mutant (LpΔ*dotA*) at an MOI of 10 or left uninfected but treated with LPS (100 ng/ml). All cells were treated with 1000 pg/ml IFN-β. The production of NO was measured via the Griess assay at the indicated times, and the data points from independent experiments and the averages of three technical replicates. The data is expressed as fold change compared to uninfected/treated cells. A two-way ANOVA with Geisser-Greenhouse correction and Tukey’s multiple comparisons test was performed on all data. Significance is indicated as follows: \**p* ≤ 0.05, \*\**p* ≤ 0.01, and \*\*\*\**p* ≤ 0.0001.

Next, the role of the T4SS on the capacity of infected cells to produce NO upon IFN-α stimulation was analyzed. Uninfected RAW-Lucia ISG-KO-IRF3 cells treated with 100 ng/ml LPS and 1000 pg/ml IFN-β were used as a positive control. RAW-Lucia ISG-KO-IRF3 cells were infected with either wild-type *L. pneumophila* or Δdot*A* mutant *L. pneumophila* at an MOI of 10 and treated with IFN-β (1000 pg/ml). The wild-type *L. pneumophila*-infected cells had significantly less NO production than the Δdot*A* mutant-infected cells or the positive control cells (Figure 4B). The level of NO produced was not significantly different between the Δ*dotA* mutant *L. pneumophila*-infected cells and the positive control. Overall, these results demonstrate that the Dot/Icm T4SS is necessary for the bacteria to inhibit IFN-β-mediated cell signaling and suggest that one or more *L. pneumophila* effectors secreted into the host cell are responsible for inhibiting the IFN-β signaling pathway in murine cells.

To explore if the IFN-β signaling inhibition caused by *L. pneumophila* seen in the murine cells also occurs in human cells, the human monocyte cell line THP-1 was used. THP-1 cells were differentiated into macrophages using PMA for 48 hours and then cultured in media without PMA for 48 hours, before infection with bacteria. The effects of *L. pneumophila* on the IFN-β signaling pathway were observed by flow cytometry measuring the cell surface tetherin levels. For these experiments, GFP wild-type and GFP Δ*dotA* mutant *L. pneumophila* were used, which allowed us to specifically study *L. pneumophila*-infected THP-1 cells. The gating strategy for the flow cytometry analysis was first to define the cell population (SSC/FSC), then gate on single-cells, live cells, and finally the GFP-positive, infected cells (Supplementary Figure 2). The tetherin antibody showed an increase in mean fluorescence intensity (MFI) when compared with the isotype control (Supplementary Figure 3A), and the GFP signal did not influence the APC tetherin antibody signal (Supplementary Figure 3B). Differentiated THP-1 cells were infected with wild-type or Δ*dotA* mutant *L. pneumophila* at an MOI of 10. Then, the infected and uninfected cells were left untreated or treated with IFN-β (10 ng/ml). After 20 hours, the uninfected cells treated with IFN-β had roughly a three-fold increase in tetherin-staining MFI compared with the uninfected, untreated cells (Figure 5A). The wild-type *L. pneumophila*-infected, IFN-β treated cells had less of an induction of tetherin staining (1.5-fold over uninfected, untreated) compared to the three-fold induction of the uninfected, IFN-β-treated cells. However, the Δ*dotA* mutant *L. pneumophila*-infected, IFN-β treated cells had no significant difference in the tetherin cell surface expression when compared with the uninfected, IFN-β-treated cells (Figure 5A). These results show that *L. pneumophila* inhibits the IFN-β signaling pathway in a T4SS-dependent manner in human macrophage-like cells.

**Figure 5:**
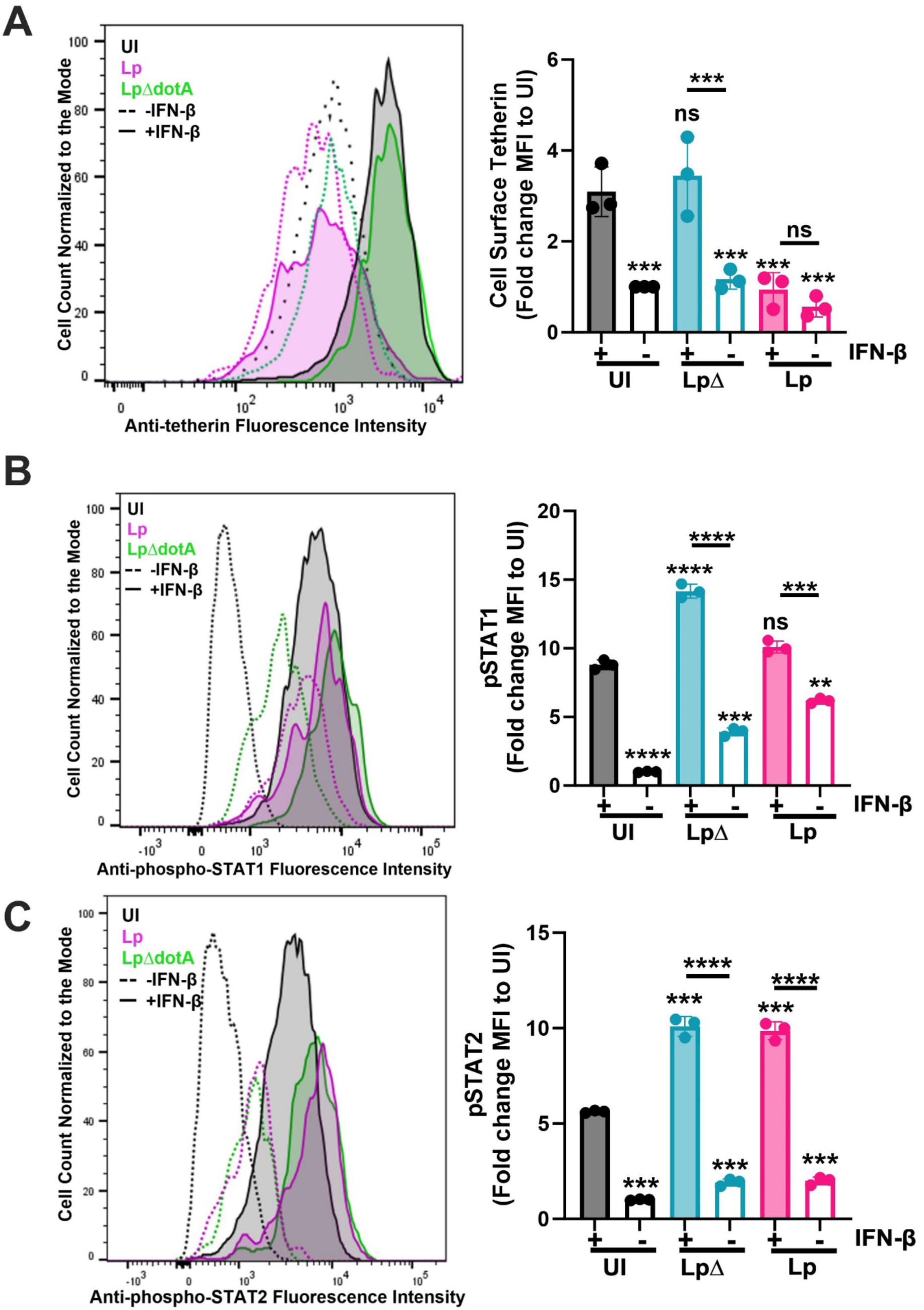
*L. pneumophila* inhibits cell surface expression of tetherin but does not affect the phosphorylation of STAT1 and STAT2. THP-1 cells were infected with GFP-expressing wild-type *L. pneumophila* (Lp) and *dotA* deficient mutant (LpΔ) at an MOI of 10 and treated or not with 10 ng/ml IFN-β. (A) The flow cytometry analysis performed on the GFP+/infected cells using a fluorescent antibody targeting the cell surface ISG protein tetherin at 20 h.p.i.. (B) The flow cytometry analysis performed on the GFP+/infected cells at an MOI of 10 using a fluorescent antibody targeting phosphorylated STAT1 30 minutes post infection. (C) The flow cytometry analysis performed on the GFP+/infected cells at an MOI of 10 using a fluorescent antibody targeting phosphorylated STAT2 30 minutes post infection. The histograms are from one experiment representative of three independent experiments. The bar graphs show the mean fluorescent intensity (MFI) fold changes compared with the uninfected untreated cells for each flow cytometry analysis. Individual points represent independent experiments. An asterisk indicates multiple unpaired *t-*tests with Welch correction relative to the uninfected IFN-β treated cells. Significance is indicated as follows: \*\**p* ≤ 0.01, \*\*\**p* ≤ 0.001, and \*\*\*\**p* ≤ 0.0001.

The activation of STAT1 and STAT2 via phosphorylation after IFN-β binding to IFNAR1 is an important step in the IFNAR signaling pathway and is important for the suppression of *L. pneumophila* growth (Abdul-Sater et al., 2015). In order to assess if *L. pneumophila* infections inhibit STAT1 or STAT2 phosphorylation, the levels of phospho-STAT1 and phospho-STAT2 in *L. pneumophila-*infected THP-1 cells treated with IFN-β were measured by flow cytometry. The total STAT1, phospho-STAT1, total STAT2, and phospho-STAT2 specific antibodies showed good increases in MFI when compared with the isotype controls (Supplementary Figure 4), and for each of these antibody stainings, the GFP signal of the *L. pneumophila* bacteria did not cross over (Supplementary Figure 4). Differentiated THP-1 cells were infected with wild-type or Δ*dotA* mutant *L. pneumophila* at an MOI of 10. Next, the infected and uninfected cells were left untreated or treated with IFN-β (10 ng/ml) for 30 minutes. The uninfected, IFN-β-treated cells had about a nine-fold increase in phospho-STAT1 MFI and a six-fold increase in phospho-STAT2 MFI when compared with the uninfected, untreated cells, indicating that IFN-β increased the phosphorylation of these two proteins (Figures 5B and 5C). The wild-type *L. pneumophila* and Δ*dotA* mutant *L. pneumophila* infected, untreated cells had increased phospho-STAT1 (Lp: 6-fold change, LpΔ: 3-fold change) and phospho-STAT2 (Lp: 2-fold change, LpΔ: 2-fold change) levels compared with the uninfected, untreated cells, probably due to endogenous expression of IFN-β after the infection (Figures 5B and 5C). Cells infected with wild-type *L. pneumophila* and treated with IFN-β had a similar fold change increase in STAT1 phosphorylation compared to uninfected, IFN-β-treated cells (about 9 to 10-fold for both). However, these cells showed a significantly stronger increase (10-fold) in the level of phospho-STAT2 when compared with the uninfected, IFN-β-treated cells (5-fold) (Figures 5B and 5C). Cells infected with the Δ*dotA* mutant and treated with IFN-β had significantly increased phospho-STAT1 and phospho-STAT2 levels compared with the uninfected, IFN-β treated cells (Figures 5B and 5C). The levels of total STAT1 and STAT2 were significantly changed in some conditions but overall, the magnitude of these changes were minor (Supplementary Figure 5A and B). These results suggest that the IFN-β signaling pathway inhibition caused by *L. pneumophila* is downstream of the phosphorylation of STAT1 and STAT2.

## Discussion

Type I IFNs have mainly been shown to play a protective role during viral infections (Lazear et al., 2019); however, the role of type I IFNs is more complex in bacterial infections since IFN-β has been shown to produce protective and detrimental effects depending on the bacterial pathogen (Boxx and Cheng, 2016; Peignier and Parker, 2021). We demonstrate that IFN-β plays a protective role against an *L. pneumophila* infection and contributes to the clearance of *L. pneumophila* in BMDMs (Fig. 3). This observation aligns with previous studies, which have shown that type I IFN protects against *L. pneumophila* infections in an IFN-γ-independent way during *ex vivo* and *in vivo* infections (Schiavoni et al., 2004; Plumlee et al., 2009; Lippmann et al., 2011; Abdul-Sater et al., 2015). NOS2/iNOS generates NO and subsequently reactive nitrogen intermediates, which have cell-intrinsic, anti-bacterial activity against the growth of many bacterial pathogens but seem to be mostly ineffective against *L. pneumophila* (Gebran et al., 1994; Akamine et al., 2007; Plumlee et al., 2009; Naujoks et al., 2016; Price et al., 2019), Accordingly, we did not observe any difference in bacterial growth between the *Nos2^-/-^*BMDMs and the wild-type BMDMs treated with IFN-β. One possibility for this lack of difference in bacterial survival could be redundancy in signaling pathways similar to what has been shown for the protective activity of IFN-γ against *L. pneumophila,* where IRG1 and NOS2 compensate for the loss of one another (Price et al., 2019). The IFN-β/IRG-1 pathway mediates increased itaconic acid production, which targets *L. pneumophila-*containing vacuoles and acts as a bactericide (Naujoks et al., 2016). Another possibility is that, unlike IFN-γ, the anti-bacterial effect of IFN-β is RNS independent.

Previous studies have identified bacterial pathogens that inhibit IFN-β signaling. For instance, *M. tuberculosis* inhibits IFN-β signaling by limiting the activation of IFNAR-associated tyrosine kinase (Banks et al., 2019a). *S. sonnei* inhibits IFN signaling through the secreted effector OspC, which binds to the host cell calmodulin, inhibiting calmodulin kinase II and STAT1 phosphorylation (Alphonse et al., 2022). *L. pneumophila* inhibits host cell IFN-β production via the T4SS-secreted effector Sdha (Monroe et al., 2009). However, we, to our best knowledge, are the first to demonstrate that *L. pneumophila* employs mechanisms to inhibit IFN-β signaling. Our results suggest that *L. pneumophila* secretes an effector(s) to inhibit the IFN-β response since the T4SS-deficient mutant failed to inhibit IFN-β signaling (Figs. 4A, 5A). Furthermore, *L. pneumophila* inhibits IFN-β signaling downstream of STAT1 and STAT2 phosphorylation since they are not downregulated upon *L. pneumophila* infection (Figs. 5B, 5C), which is different from *M. tuberculosis*- and *S. sonnei*-mediated inhibition (Banks et al., 2019b; Alphonse et al., 2022).

Humans are an accidental host for *L. pneumophila,* and so *L. pneumophila* has primarily adapted to survive in amoebas and not in human macrophages. Amoebas do not have an equivalent receptor to IFNAR or any class of receptors associated with tyrosine kinases (Goldberg et al., 2006). The evolutionary advantage for *L. pneumophila* to inhibit IFNAR-signaling when amoebas lack equivalent receptors is thus unclear. However, amoebas, such as *Dictyostelium sp.* and *Acanthamoeba sp.,* express a homolog to mammalian STAT proteins (Kawata et al., 1997; Kicinska et al., 2014). In particular, the amoebal STAT protein is closely related to the human and mouse STAT2 protein (Kicinska et al., 2014), which is the STAT isoform involved in type I but not type II IFN signaling (Villarino et al., 2020). This could potentially explain why *L. pneumophila* can inhibit type I but not type II IFN signaling.

SH2 domains are the primary protein domain responsible for binding tyrosine-phosphorylated targets involved in tyrosine kinase signaling cascades, such as the STAT1 and STAT2 cascade (Schlessinger and Lemmon, 2003). Interestingly, *L. pneumophila* encodes for 84 proteins that contain an SH2 domain, and these SH2 domains, while lacking pocket specificity, bind mammalian phosphorylated tyrosines with strong affinities (Kaneko et al., 2018). Consequently, *L. pneumophila* could have evolved mechanisms to suppress amoebal STAT-mediated transcriptional regulation, allowing for the growth of *L. pneumophila* in amoebas, and serendipitously, this mechanism also inhibits mammalian STAT2-mediated transcriptional regulation. Due to the homology of mammalian STAT2 with the amoeba STAT and the SH2 domains found in *L. pneumophila* proteins (Kicinska et al., 2014), we hypothesize that IFNAR signaling could be inhibited at the STAT2 level after the phosphorylation of STAT2 via interactions with an SH2 domain found in an *L. pneumophila* protein. This could potentially block the association of STAT2 with STAT1 and IRF9, which is required to form the active ISGF3 complex. However, further studies are needed to test this hypothesis and identify the putative *L. pneumophila*-secreted effector responsible for the inhibition of IFN-β signaling.

## Acknowledgements

C.N.S.A., D.A.B., M.S. and V.B. were supported by grants R21AI142396 and R21 AI151973. S.A.V. by grant R01 AI123371. M.S. was supported by “The Stefanie & Richard Vogel Graduate Student Award” and T32 AI089621.

---

**Supplementary Figure 1:**
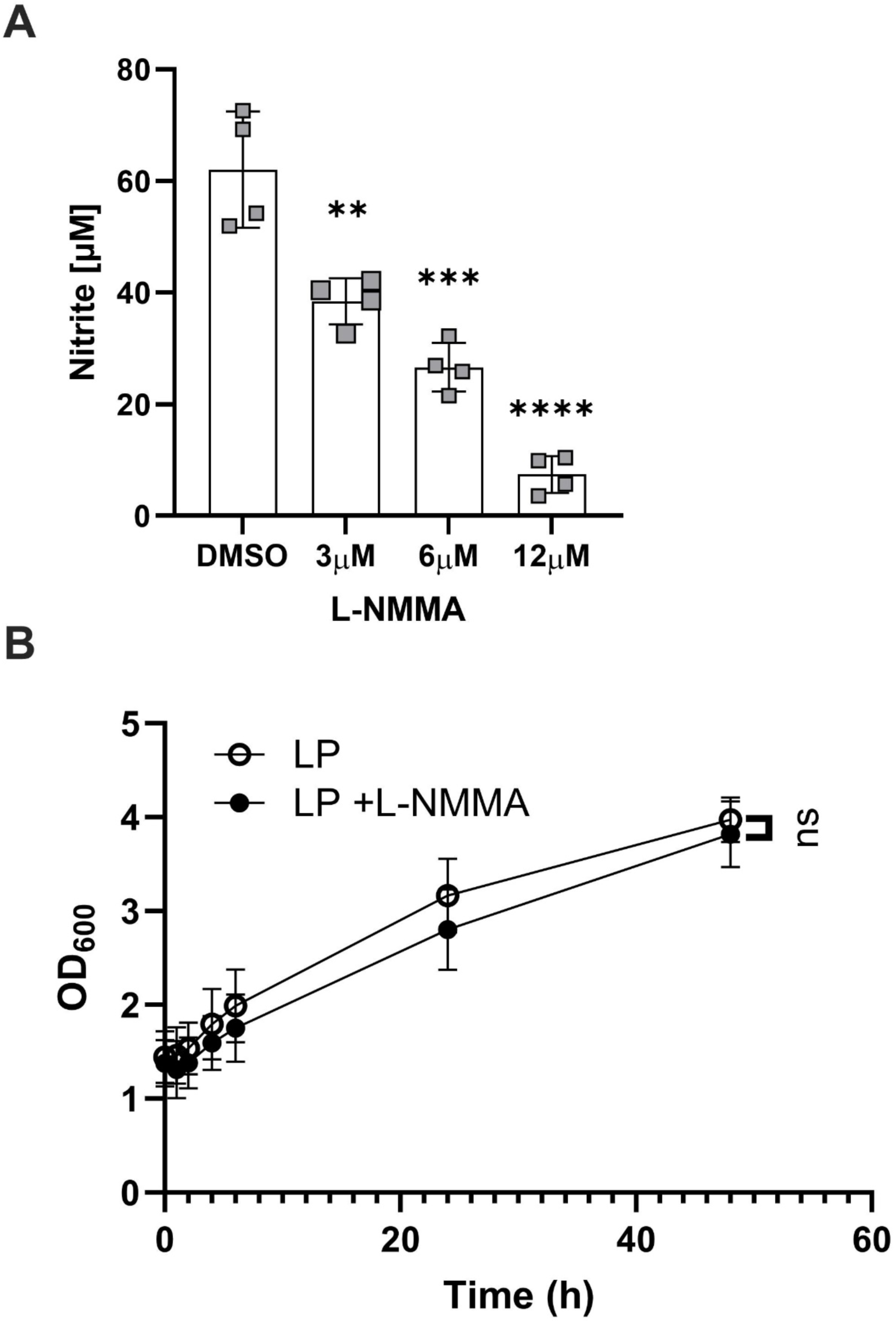
Effects of L-NMMA on nitrite production and *L. pneumophila* growth. (A) The dose-response analysis of the iNOS2 inhibitor L-NMMA on RAW-ISG-KO-IRF3 cells treated with 1000 pg/ml of IFN-β 20 hours after dosing. The data points are from four independent experiments. The statistical analysis was done using a repeat-measure (RM) one-way ANOVA. (B) The growth curve of wild-type *L. pneumophila* (Lp) measured by OD_600_ in the presence or absence of 12 µM L-NMMA. Points are the averages mean ± standard deviation of three independent experiments. The statistical analysis was done by a multiple unpaired *t-*test with Welch correction comparing the two groups at the time points indicated. Significance is indicated as follows: \*\**p* ≤ 0.01, \*\*\**p* ≤ 0.001, and \*\*\*\**p* ≤ 0.0001.

**Supplementary Figure 2:**
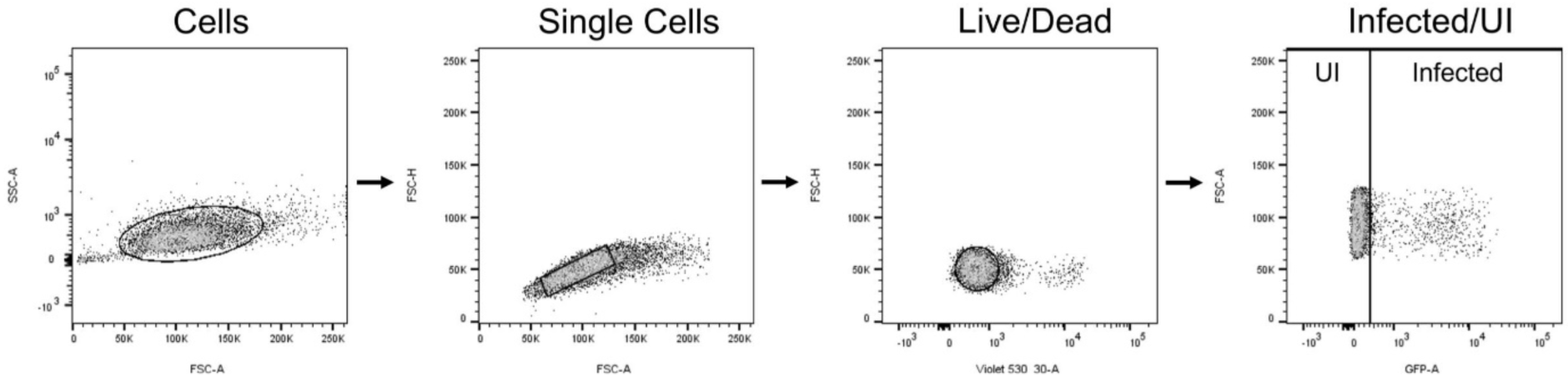
Flow Cytometry Gating Strategy. THP-1 cells were first gated to isolate cells from debris (SSC/FSC). Next single cells were selected (FSC-H/FSC-A) and using LIVE/DEAD Fixable Dye the live cells were selected. Lastly, infected cells were selected by measuring the GFP fluorescence intensity.

**Supplementary Figure 3:**
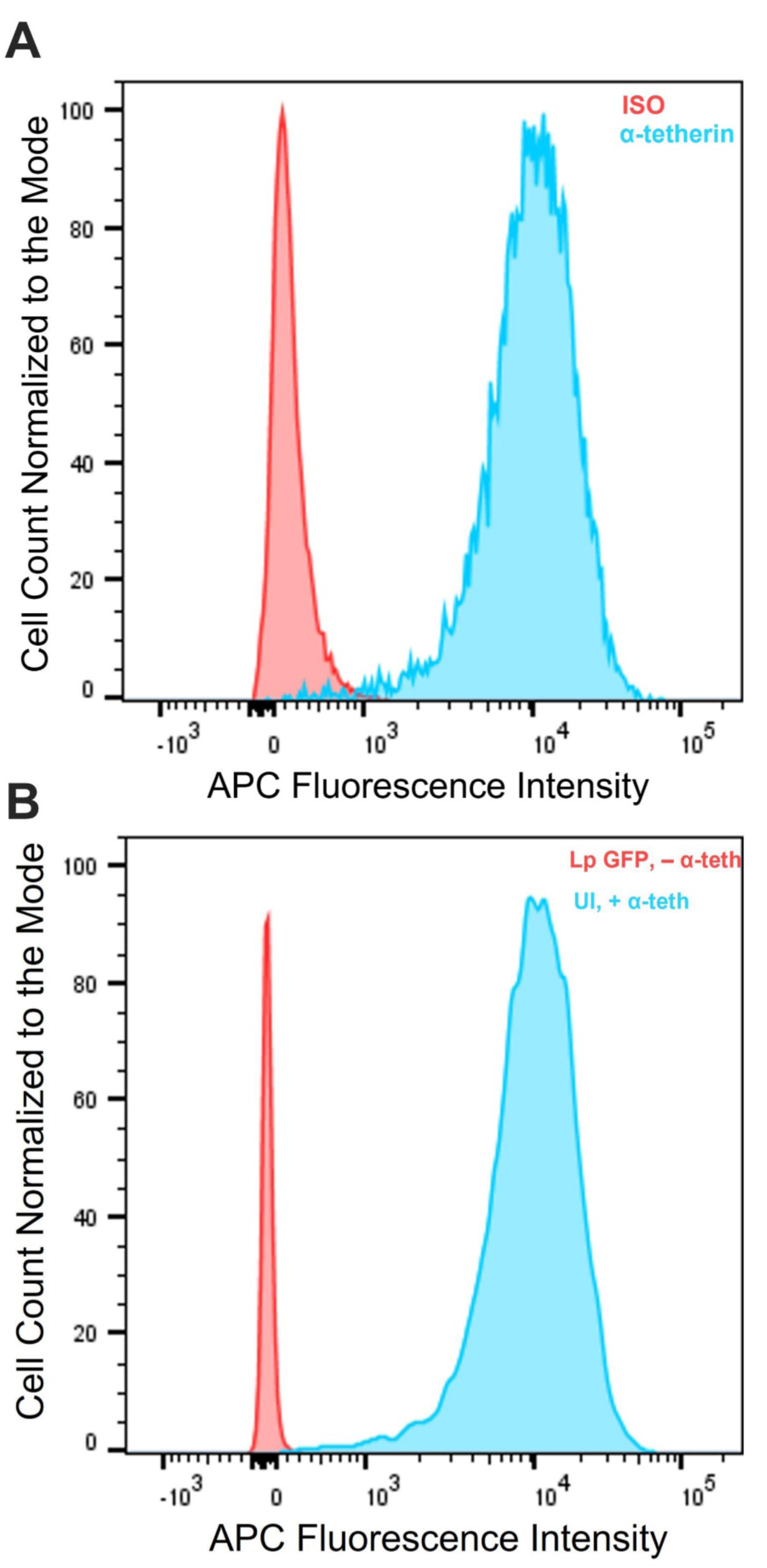
Tetherin antibody testing and GFP Flow Cytometry. (A) MFI peaks of uninfected IFN-β (10 ng/ml) treated THP-1 cells stained with the tetherin (α-tetherin) or Isotype control antibodies (ISO). (B) Histograms of uninfected THP-1 cells stained with the APC conjugated tetherin antibody (UI+IFN-α) and *L. pneumophila* GFP infected (GFP) (MOI of 10) unstained cells measuring the APC intensity.

**Supplementary Figure 4:**
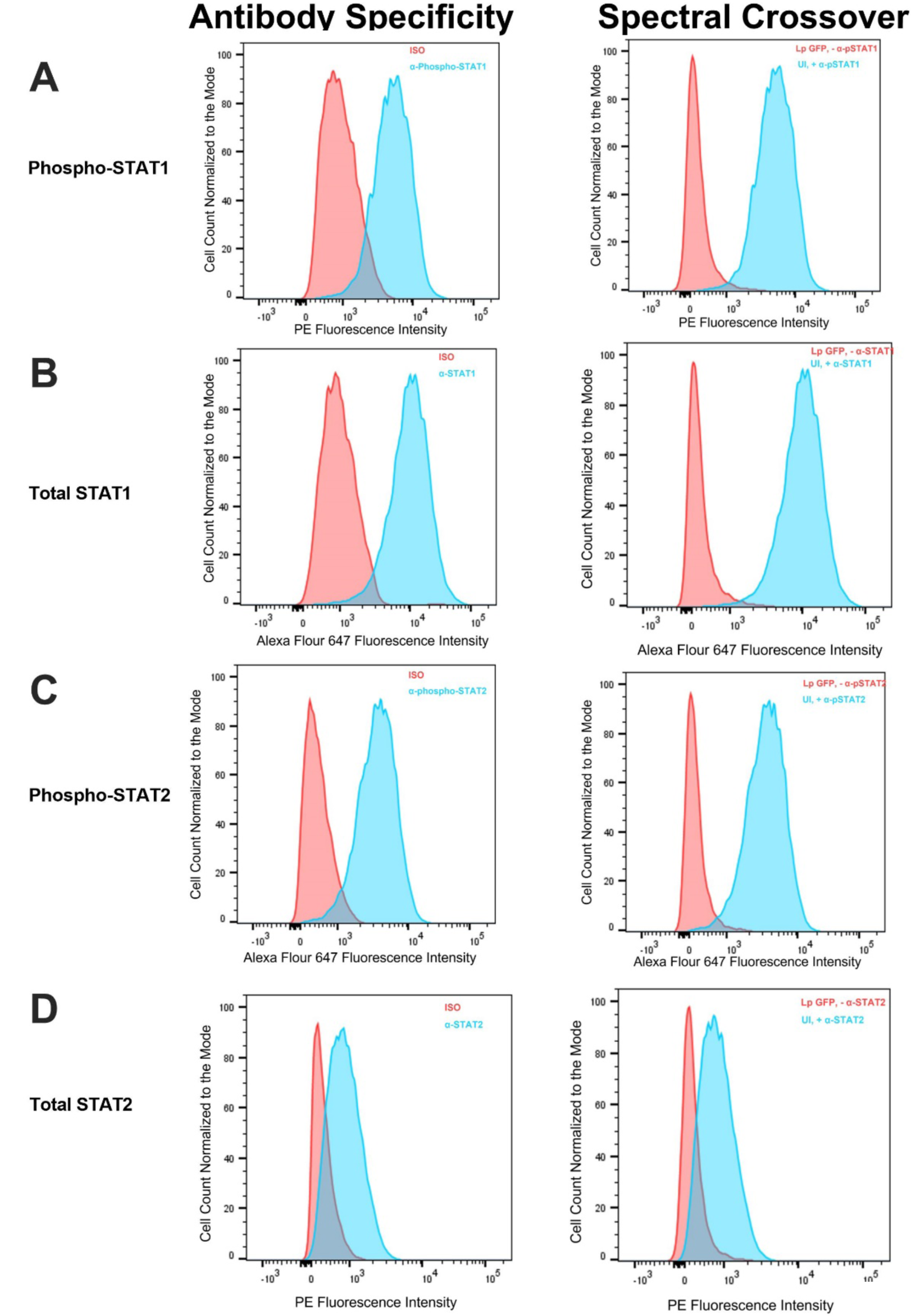
Isotype antibodies and GFP Flow Cytometry control experiments. The left column indicates the tests of the specific antibody stainings compared to isotype controls as indicated and uninfected/IFN-β-treated (10 ng/ml) THP1 cells. The right column analyses the potential spectral crossover of the GFP-signal into any of the other channels used for the specific antibody stainings by infecting cells with Lp-GFP (MOI of 10) and no specific antibody or having uninfected/IFN-β-treated cells stained with specific antibody. (A) phospho-STAT1 antibody (B) total STAT1 antibody, (C) phospho-STAT2 antibody and (D) total STAT2 antibody.

**Supplementary Figure 5:**
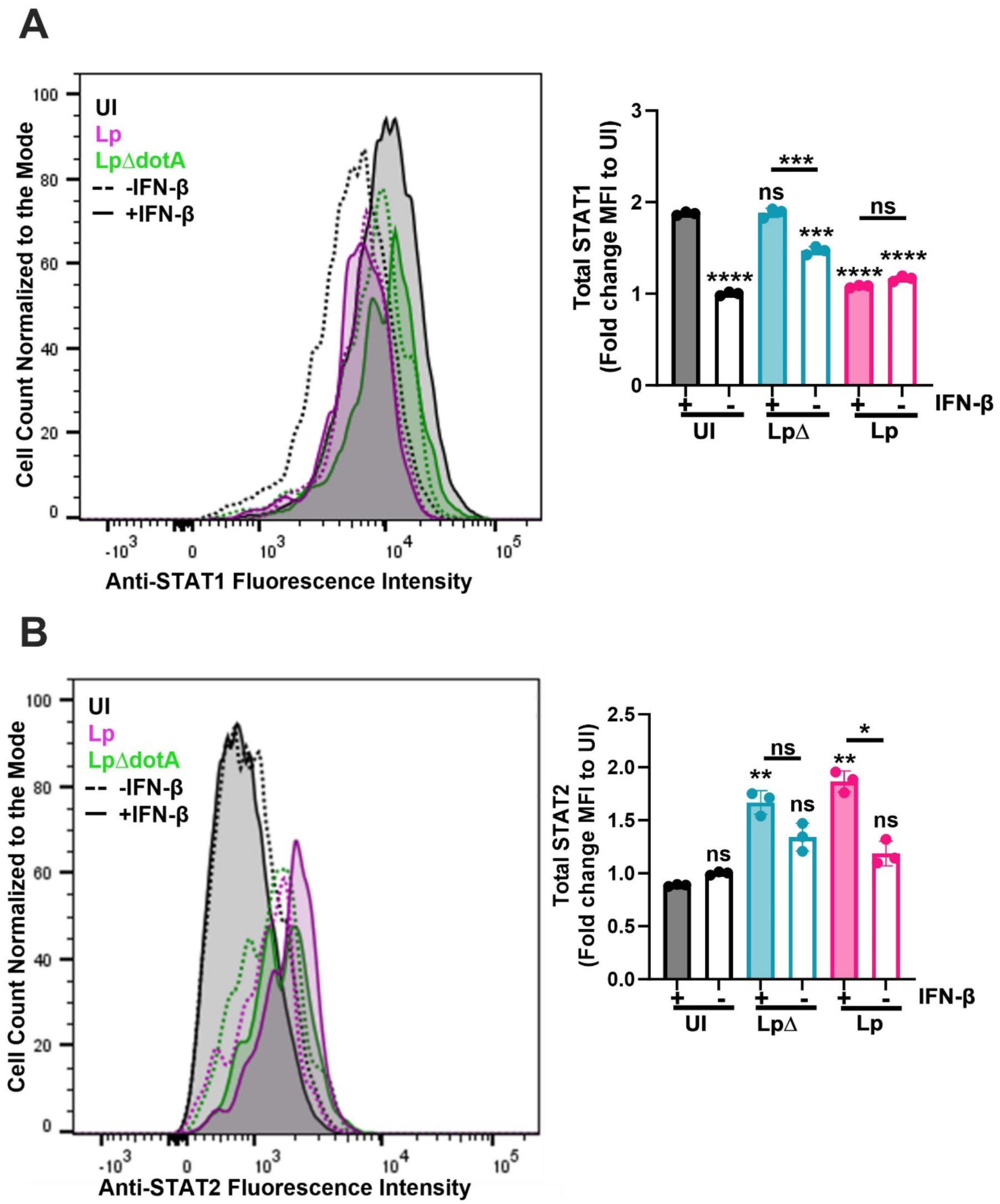
The effects of *L. pneumophila* infection on the total STAT1 and STAT2 levels. THP-1 cells were infected with GFP-expressing wild-type *L. pneumophila* (Lp) and *dotA* deficient mutant (LpΔ) at an MOI of 10. The treated groups were treated with 10 ng/ml IFN-β. (A) The flow cytometry analysis performed on the GFP+ infected cells using a fluorescent antibody targeting total STAT1 30 minutes post infection. (B) The flow cytometry analysis performed on the GFP+ infected cells using a fluorescent antibody targeting total STAT2 30 minutes post infection. The histograms are representative of three independent experiments and illustrate the mean fluorescent intensity (MFI) of each group. The bar graphs show the MFI fold changes compared with the uninfected/ untreated cells for each flow cytometry analysis. Individual points indicate independent experiments. An asterisk indicates multiple unpaired *t-* tests with Welch correction relative to the uninfected IFN-β treated cells. Significance is indicated as follows: \**p* ≤ 0.05, \*\**p* ≤ 0.01, \*\*\**p* ≤ 0.001, and \*\*\*\**p* ≤ 0.0001.

## Notes

### Competing Interest Statement

The authors have declared no competing interest.

### Summary of Updates

We have included new data addressing to more strongly support the claim that Lp inhibits IFN-beta signaling by: 1) using qRT-PCR to demonstrate that the Lp-infection inhibits IFN-beta mediated transcription, 2) showing that Lp does not induce cell death of the reporter cells and 3) using a flow cytometry-based approach to show that Lp infection inhibits the expression of the IFN-beta inducible tetherin protein on the surface of human macrophages (differentiated THP-1 cell line) in a T4SS-dependent way. in addition we provide new data showing that the IFN-beta mediated growth restriction of Lp is indeed independent of NO-production which is a change to our previous finding but in line with results from previous publications.

